# Central visceral commands formatted as the slow rhythms encoded in the temporal structures of the sympathetic firing originated from the neonatal rat spinal cord in vitro

**DOI:** 10.1101/2022.08.01.502255

**Authors:** Chun-Kuei Su, Chiu-Ming Ho

## Abstract

Central sympathetic neural circuits continuously generate efferent commands to sustain rhythmic operations of their peripheral effectors. However, beyond the firing rates, in what configuration an effective visceral command is formatted by the spiking activities was largely unknown. This study used an in vitro splanchnic nerve–spinal cord preparation of the neonatal rats as an experimental model and recorded spontaneous efferent activities from single sympathetic fibers. The patterns of the fiber activities were quantitatively evaluated by a metric, the so-called local variation (Lv). Lv was derived from calculating the relative differences between adjacent interspike intervals (ISIs), and thus, described the spiking patterns from a regular spiking with Lv = 0 to that spiking in bursts with Lv >1. Along the time course, the dynamic components of Lv (dLv) displayed quasiperiodic oscillations. Continuous wavelet analysis showed that dLv oscillations registered a dominant power rhythm at ∼7 mHz. This slow rhythmicity was heterogeneously altered by application of various antagonists that interrupted endogenous neurotransmitter activities mediated by ionotropic glutamate receptors or GABA_A_ receptors in the spinal cord. Thus, the oscillation of dLv manifested itself as a feature of neural network operation. On the assumption that the total power of the dLv oscillations reflects an activity status of neural network operation, the antagonist-induced change of the power fits well with a concomitant ISI change in a negative sigmoid relationship, which explains the heterogeneity of the antagonist-induced firing responses. In conclusion, dLv oscillations reflect a dynamic status of neural network operation. The slow rhythms embedded in the dLv oscillations are likely acting as an information coder and convey effective central visceral commands that can be followed by their downstream effectors.

**Author summary:** In what format the information is encoded by the neural activity is relatively unknown. This is especially true for autonomic regulation of visceral functions. We seek to determine the format of the central sympathetic commands, by which it can generate an effective driving force to regulate the operation of their peripheral target organs. Because most visceral organs operate with certain rhythms, we anticipate that the central visceral commands are also formatted in rhythms. Two aspects of techniques were employed. One was the in vitro electrophysiological and pharmacological techniques to acquire and to manipulate the neural signals recorded from sympathetic single-fibers. The other was the computational techniques to extract the information embedded in the timing of the spiking behaviors and to examine how these features were altered by pharmacological manipulations. We found a spontaneous change of sympathetic spiking patterns displaying rhythms with a frequency that could match the rhythmic operation of many visceral organs. Thus, the meaningful information carried by the sympathetic neural circuits for the visceral controls is likely to be encoded in the dynamics of their spiking patterns. In summary, a dynamic change of neural spiking patterns could be used as a simple scheme for neural information coding.

## Introduction

Information encoded in a neural signal is not only conveyed by the appearance of neural discharges but also by the temporal structures built between the neural discharges. In terms of single neural activity, the temporal structures are constructed by a series of interspike intervals (ISIs) with orderly arrangements. The inverse of the ISI being counted as the firing rate is well known as a neural representation of intensity. Less well understood is the temporal order of ISIs, namely, the ISI sequence representing for in neural signal processing. Unique features encoded in the ISI sequences have been implicated to carry information of visual stimuli [1–3], display activity history in pacemaker neurons [4], and differentially affect the transmission efficiency of individual synapses for selective communication between neurons [5]. In the aspect of the efferent signals generated from the mammalian central nervous system, how the ISI sequences participate in constructing an effective visceral command is largely unknown.

Sympathetic nerve discharges (SND) wax and wane over time, forming an envelope activity of sinusoids, which likely represents a pattern of information coding at the population level. Analysis of this sinusoidal activity reveals the rhythms of SND in ranges from tenth of Hz up to 10 Hz in human and other animals [6–10], and ∼0.5‒2 Hz in a reduced brainstem‒spinal cord preparation of neonatal rats [11]. While it is generally believed that a coherence of ensemble neural activity underlies the rhythmogenesis [12, 13], the functional significance of the SND rhythms remains elusive [14]. In a close-loop control circuit, an efferent command generated from the neural circuits produces an effect that may subsequently feedback to the loop by producing a delayed afferent signal. For instance, an increased arterial blood pressure resulted from increased efferent sympathetic commands would stimulate the baroreceptors, and with a latency, the baroreceptor-mediated inhibitory afferent signal would feed into the operation of the central neural circuits and subsequently affect the next episode of the efferent command generation. As a physiological or developmental adaptation, a well-predicted response time could be blueprinted in the central neural circuit, thereby leading to an inherent waning following a waxed command signal. Operation of such a neural scheme that encodes the hardwire design for the response time is likely manifested by the temporal order of the neural commands, and being observed as the neural rhythms at the population level.

The cardiovascular system is one of the effectors controlled by the sympathetic nervous system. While the sympathetic nerves convey impulses of the rhythms usually locked in several Hz, classical studies using electrical stimulation of sympathetic nerves have demonstrated that sympathetic effectors are low-pass filters and can only faithfully respond to the stimulus frequency at <0.02 Hz [15–18]. The mismatch in the rhythmic frequency range between the commanding neural signals at several Hz and the resonant response of their effectors at hundredth of Hz argued against the SND rhythms as a direct information coder. As this frequency mismatch was barely explained in the past studies, the current work aimed to elucidate if the central sympathetic neural networks would generate a neural code in formats of rhythms that could be faithfully translated by their effectors.

The goal of this study is to elucidate if working information is configured by the ISI sequences. As the ISI sequences describe the spiking patterns, a quantitative description of the irregularity of the spiking patterns requires a metric to evaluate ISI variations. In the past studies, different metrics have been used, including coefficient of variation (Cv), local variation (Lv), and etc. [19–21]. For simplicity, Lv was chosen as the main metric to describe the spiking irregularity here for two reasons [22]. First, Lv is an index of spiking patterns that may change from a regular firing of constant ISIs with Lv = 0 to a firing in irregular bursts with Lv > 1; when Lv > 1, the efficiency of synaptic transmission is enhanced by burst firing. When Lv = 1, the spiking patterns are constructed by a series of ISI with the Poisson probability distribution; i.e., given a fixed value of the ISI means, the spikes occur randomly. Second, compared to Cv, Lv is relatively independent from the influence of the firing rates or the ISI contents themselves, because its calculation only takes into account of the differences between neighboring ISIs. Thus, Lv was chosen as an ideal metric to evaluate if information coding could be embedded in the ISI sequence.

Single-fiber activities spontaneously generated in the in vitro thoracic spinal cord preparations of the neonatal rats were recorded from collagenase-dissociated splanchnic sympathetic nerves [23, 24]. As the dynamic components of Lv (dLv) that revealed the single-fiber spiking patterns along the time course displayed apparent oscillations, the rhythms contained in dLv oscillations were further elucidated by continuous wavelet analyses. The hypothesis to be examined here is that dLv oscillations in providing a moment-to-moment adjustment of synaptic efficiency to their downstream effectors can imprint a rhythm that is within the operating frequency range of the sympathetic effectors. Moreover, because endogenous amino acid neurotransmitter activity could reshape whole-nerve SND in tonic or bursting forms [25], whether dLv oscillations manifested themselves as a rhythmic operation of the spinal neural network would be tested by application of various antagonists that interrupted the endogenous neurotransmissions to explore the pharmacological basis of dLv rhythmogenesis.

## Results

### Effects of the drug application on the metrics describing spiking rates and variations

Analyses were based on neural recordings of 84 sympathetic single-fibers, which were spontaneously active under control conditions. The means of ISIs (mISIs) of individual single-fibers ranged from 0.3 to 38.7-s under control and from 0.3 to 42.8-s after application of L-2-amino-5-phosphonopentanoic acid (APV, an NMDA receptor blocker, *n* = 22 in 8 preparations), 6-cyano-7-nitroquinoxaline-2,3-dione-disodium salt (CNQX, a non-NMDA receptor blocker, *n* = 17 in 8 preparations), kynurenic acid (KYN, a broad-spectrum ionotropic glutamate receptor blocker, *n* = 27 in 17 preparations) or picrotoxin (PIC, a GABA_A_ receptor blocker, *n* = 18 in 7 preparations). Fig 1 summarizes the ISI-derived metrics prior to and after the drug tests. Noticeably, under control conditions, not all the metrics were comparable between different testing groups, indicating an apparent sampling bias between groups. Besides, applications of receptor blockers here were aimed to perturb the endogenous activity of the neural circuit operating in the spinal cord rather than to compare the effects elicited by different drugs. For simplicity, we only evaluated the drug effects within the same testing groups.

**Fig 1.**
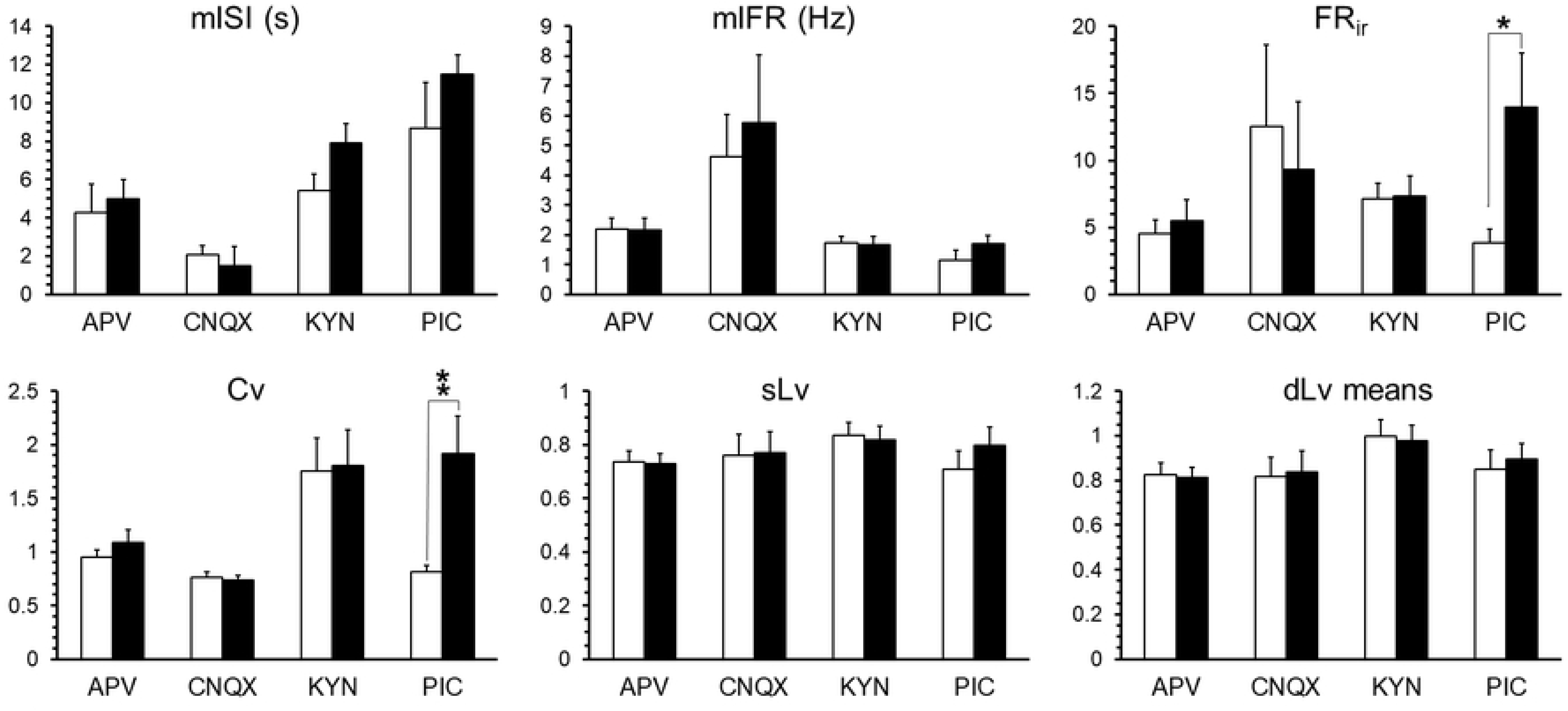
Summary of the metrics describing the spiking rates (i.e., mISI and mIFR) and the spiking variations (i.e., FR_ir_, Cv, sLv, and dLv means). Open bars, the metric values under control; solid bars, the metric values after antagonist application. All values represent the populations means of the metric values acquired from individual fiber activities (*n* = 22 in APV, 17 in CNQX, 27 in KYN, and 18 in PIC). Significance of the antagonist-induced changes from controls was verified by Student’s t-test: *, *P* < 0.05; **, *P* < 0.005. Application of PIC significantly increased FR_ir_ (control vs. PIC application: 3.87 ± 1.02 vs. 13.99 ± 3.98) and Cv (control vs. PIC application: 0.81 ± 0.06 vs. 1.91 ± 0.35).

The response of sympathetic firing to an antagonist application is heterogeneous and largely depends on its basal activity, i.e., application of the same antagonist tends to suppress the firing in the fibers with higher basal activity and enhance the firing in the fibers with lower basal activity [23, 26–28]. Due to the heterogeneity in the drug responses, Fig 1 shows that application of APV, CNQX, KYN or PIC does not significantly change the population means of mISI or the population means of the means of instantaneous firing rates (mIFR). Herein, we focused on examining the antagonistic effects on the spiking patterns both at the population level and in individual fibers. At the population level, application of APV, CNQX or KYN did not significantly change the means of Cv, firing rate irregularity (FR_ir_), dLv means (i.e., the means of a series of dLv acquired under an experimental condition) or static Lv (sLv, a metric describing a general spiking pattern under an experimental condition). In contrast, application of PIC significantly increased the means of Cv and FR_ir_, suggesting a relatively dominant role of endogenous GABA_A_ receptor-mediated activity in shaping the spiking variations at the population level. Application of PIC, however, did not significantly change the population means of sLv or dLv means, indicating that the sLv or the dLv means were statistically less sensitive than the Cv or the FR_ir_ in detecting a drug-induced change in the spiking variations at the population level.

Firing responses in individual fibers were further examined. Fig 2 shows the time course of the oscillatory patterns of IFR and dLv in 3 fiber activities that have sLv values ranging from 0.12 to 1.4, depicting a correlation of sLv values with distinct spiking patterns that varied from regular to bursty. Fig 2 also shows a change of dLv oscillatory patterns along the time course of the drug application, leading to a substantial change of sLv in individual fibers. Thus, differing from the seemingly ineffective drug effects at the population level in which the data from different fibers were pooled, the drug effects on spiking patterns were apparent in individual fibers. Statistical evaluation of the drug effects on changing sympathetic spiking patterns in individual fibers was conducted by comparing the dLv means under control conditions with the dLv means after an antagonist application. Student’s *t*-test confirmed a significant drug-induced change in the dLv means in 19 / 22 fibers by APV application, 15 / 17 fibers by CNQX application, 21 / 27 fibers by KYN application, and 13 / 18 fibers by PIC application. Taken together, the dLv means in 68 / 84 fibers were altered by the drug applications. Using the dLv sensitivity to drug applications as the categorical criteria, the metrics observed under control conditions were compared between the sensitive group (n = 68) and the insensitive group (n = 16). Significant differences were observed in the population means of mISI (sensitive vs. insensitive group in s: 4.05 ± 0.56 vs. 9.81 ± 2.78, *P* < 0.05), sLv (sensitive vs. insensitive group: 0.74 ± 0.03 vs. 0.90 ± 0.03, *P* < 0.001), and dLv means (sensitive vs. insensitive group: 0.83 ± 0.04 vs. 1.10 ± 0.06, *P* < 0.001). Thus, under control conditions, the fibers exhibiting lower mISI, lower sLv or lower dLv means tend to be more sensitive to the drug applications and change their spiking patterns. These observations implicate that the heterogeneous response to the drug applications is rooted in the basal spiking activity, i.e., those with higher basal spiking activity (i.e., lower mISI) and lower Lv (i.e., more regular firing) tend to be more sensitive to the drug application and have their spiking patterns changed by the drug applications.

**Fig 2.**
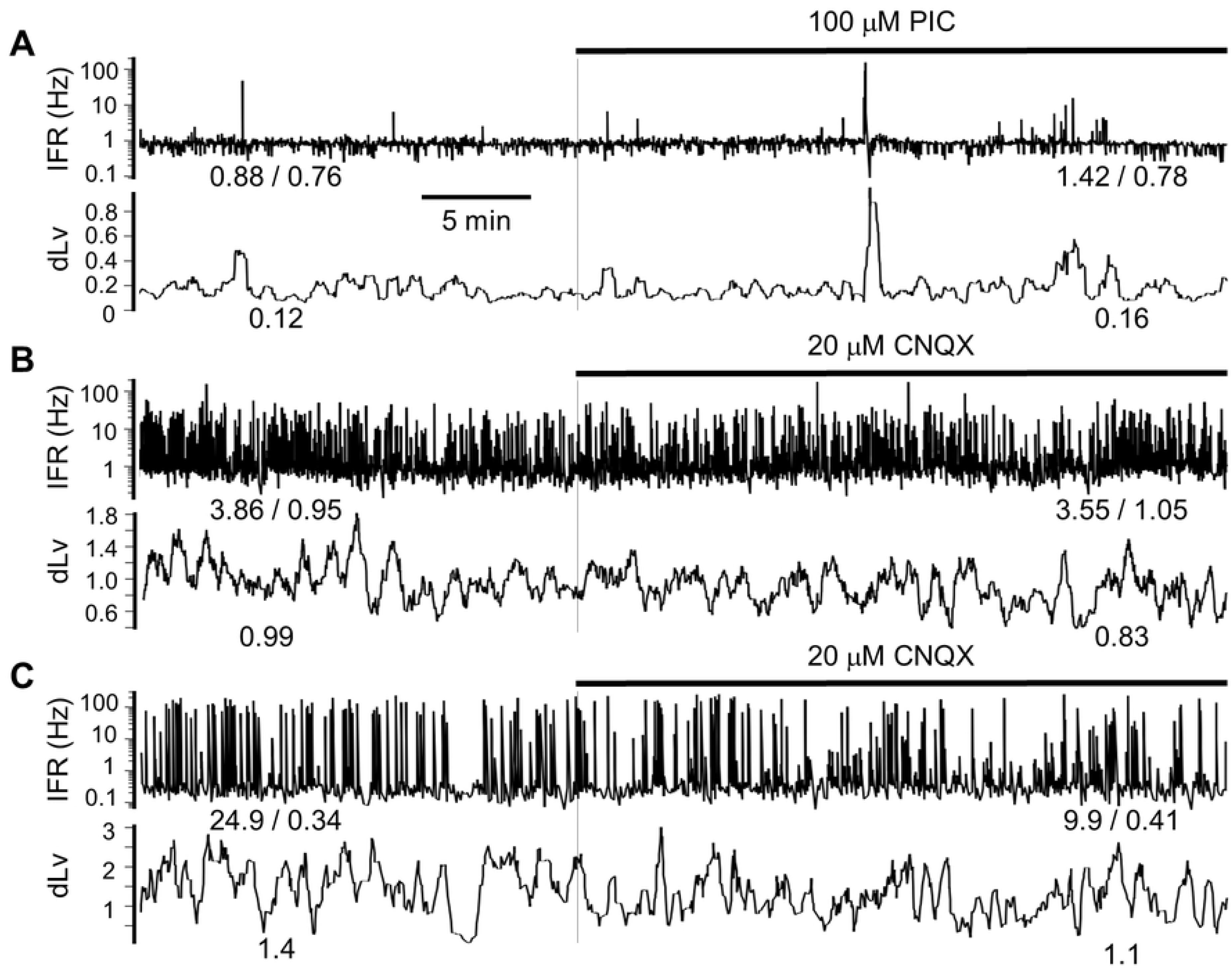
Examples showing the oscillatory patterns of IFR and dLv along the time course of experiments. Numerical values at the bottom of the traces are mIFR / AFR and sLv, respectively. From **A** to **C**, examples show an increased sLv along with an apparent increase in the irregularity of the spiking patterns. Application of the antagonist to interrupt endogenous inhibitory or excitatory neurotransmitter activity substantially altered the spiking regularity.

Because sLv is a simple metric describing the spiking patterns under an experimental condition, heterogeneity in the drug-induced change in the spiking patterns was quantitatively evaluated by analyzing the drug-induced change in sLv. Plotting the sLv after the antagonist applications versus the sLv under control conditions showed a data distribution that was well described by a power-law function (Fig 3). Little effect on sLv change was observed from application of APV, CNQX or KYN, showing a data distribution with exponential powers close to 1. In contrast, prominent effects were produced by application of PIC, showing a data distribution with an exponential power of ∼0.4. Noticeably, PIC application tends to increase sLv in those fiber activity with control sLv <0.8 and decrease sLv in those with control sLv >0.8 (Fig 3D). Using the regressed power-law equations, antagonistic effects on sLv change were simulated. Fig 4 demonstrates that application of PIC is effective to increase and decrease sLv values in those with control sLv <0.8129 and >0.8129, respectively. Thus, a PIC-sensitive endogenous neurotransmitter activity, presumably the GABA_A_ receptor activity, was involved in setting the basal spiking patterns and helped to diversify spiking patterns in a heterogeneous manner. Otherwise, the spiking patterns of different fibers would be concentrated at the one with sLv = 0.8129 (cf. an example of sLv = 0.83 was shown in Fig 2B).

**Fig 3.**
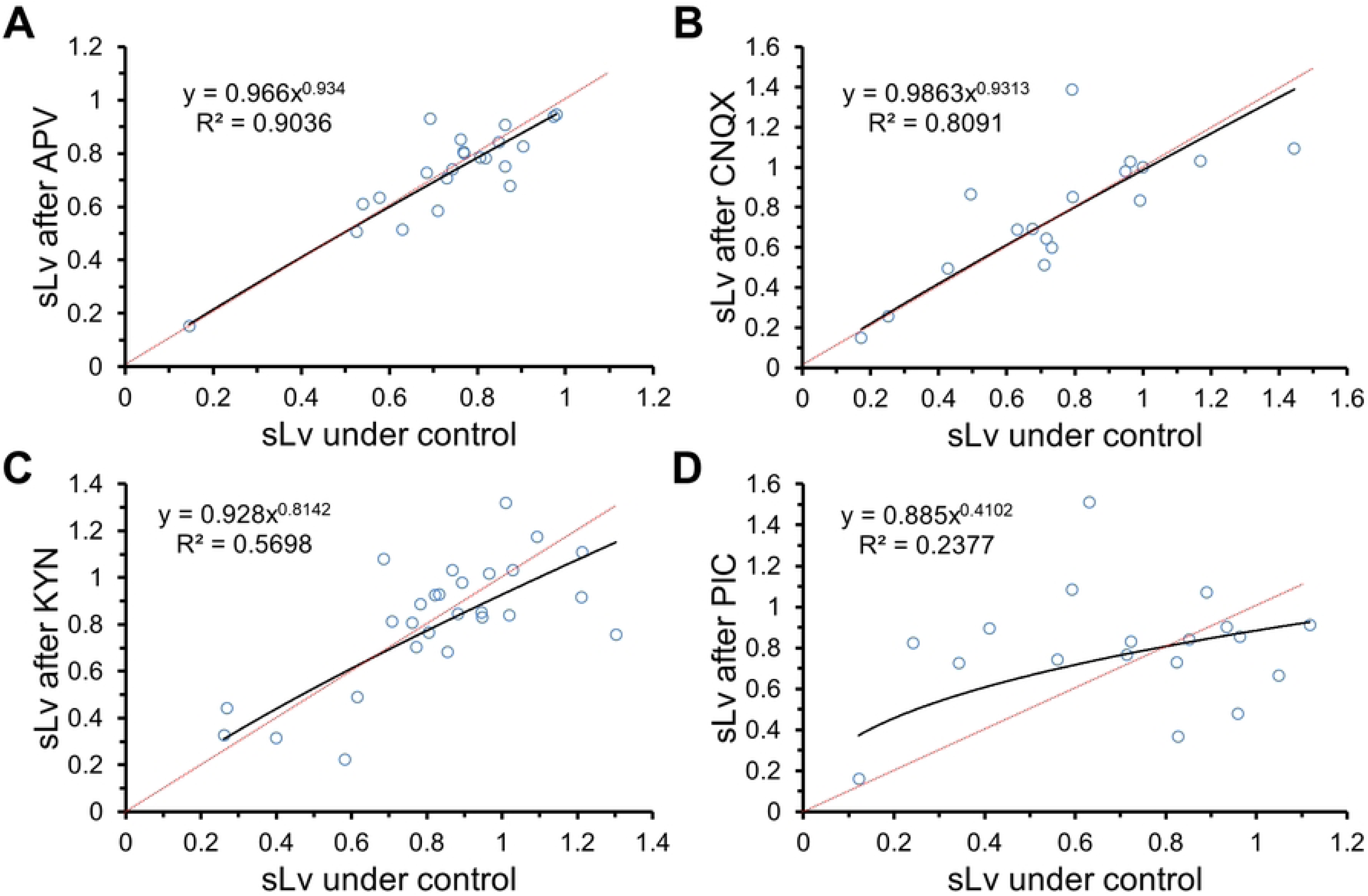
Plots of the sLv after antagonist application versus the sLv under control conditions. All the data distributions were regressed by the power law functions. Significance of the regression: *P* < 0.001 in **A‒C** and *P* < 0.05 in **D**. Red-dashed lines depict the null effects with y = x. The extents of the data distribution deviated from the red-dashed lines illustrate the antagonistic effects. Application of PIC was effective to change sLv.

**Fig 4.**
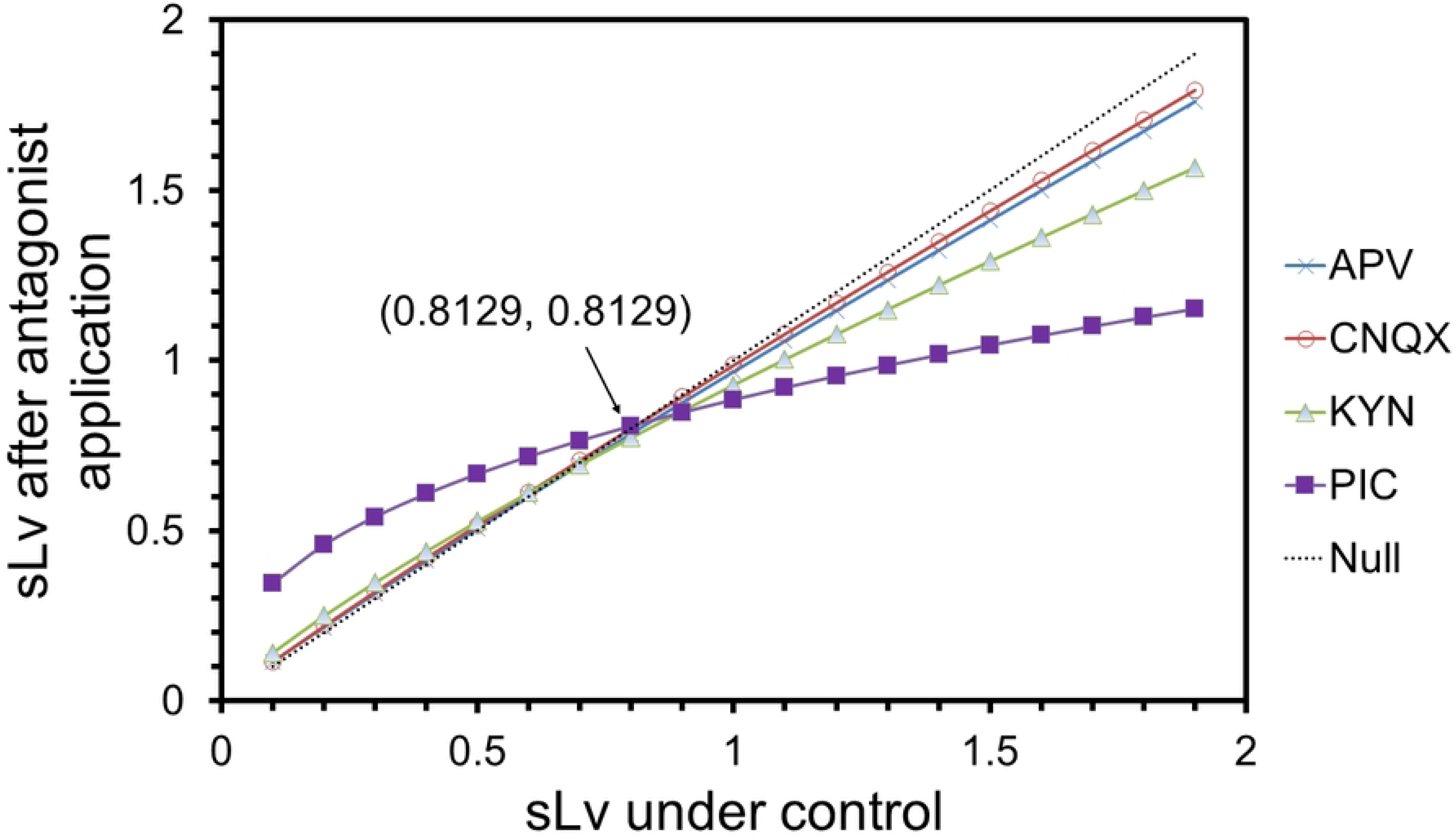
A simulated plot showing the sLv change after antagonist application. The power law functions used for simulation are acquired from the power regression lines in **Fig 3**: APV, y = 0.966x^0.934^; CNQX, y = 0.986x^0.931^; KYN, y = 0.928x^0.814^; PIC, y = 0.885x^0.410^. Dashed line shows the null effect with y = x. Deviation of the simulated curves from the dashed line illustrates the antagonistic effects. Application of PIC prominently changes the sLv, showing a decreased sLV when control sLv >0.8129 and an increased sLv when control sLv <0.8129.

### The rhythmicity encoded in dLv oscillations

Continuous wavelet analysis does not require a presumption of signal stationarity and biological signals are often not stationary. Thus, continuous wavelet analysis was selected and used to elucidate the rhythmicity encoded in dLv oscillations. Fig 5 uses the color maps to display the intensity (i.e., coefficients of wavelet transforms (Cwt)) or the energy (i.e., probability density of energy (PD_E_)) of dLv oscillations in both time- and frequency-domains, showing that dLv oscillations encode a quasiperiodicity <30 mHz and the encoded rhythms are substantially altered by application of KYN. To better reveal the rhythmicity of dLV oscillations and evaluate drug-induced change of energy profiles in the frequency-domain, epochs of PD_E_ prior to and after a drug application were averaged respectively across the time-domain and the acquired means of PD_E_ (mPD_E_) curves were compared. Fig 6 shows examples of superimposed mPD_E_ curves that were acquired under control conditions and after drug application. Drug-induced change in mPD_E_ curves was quantitatively evaluated by coefficients of deviation (C_d_). Higher C_d_ with values close to 1, as shown in Fig 6A, indicates a greater extent of dissimilarity between the control mPD_E_ curve and the mPD_E_ curve after a drug application, reflecting a stronger drug effect. In contrast, lower C_d_ with values close to 0, indicates a less dissimilarity or a higher similarity in between, reflecting a weaker drug effect (Fig 6C). Among the drug tests of 84 fibers, a wide distribution of C_d_ was observed in each testing group, suggesting a heterogeneous drug response in individual fibers. At the population level, averaged C_d_ from individual fiber responses in a testing group was 0.474 ± 0.045 in APV, 0.595 ± 0.044 in CNQX, 0.510 ± 0.042 in KYN, and 0.518 ± 0.054 in PIC. Student’s *t*-tests confirmed that each of the C_d_ means was significantly >0 with *P* < 0.0001, indicating that endogenous activities mediated by ionotropic glutamate receptors and GABA_A_ receptors involved in dLv rhythmogenesis.

**Fig 5.**
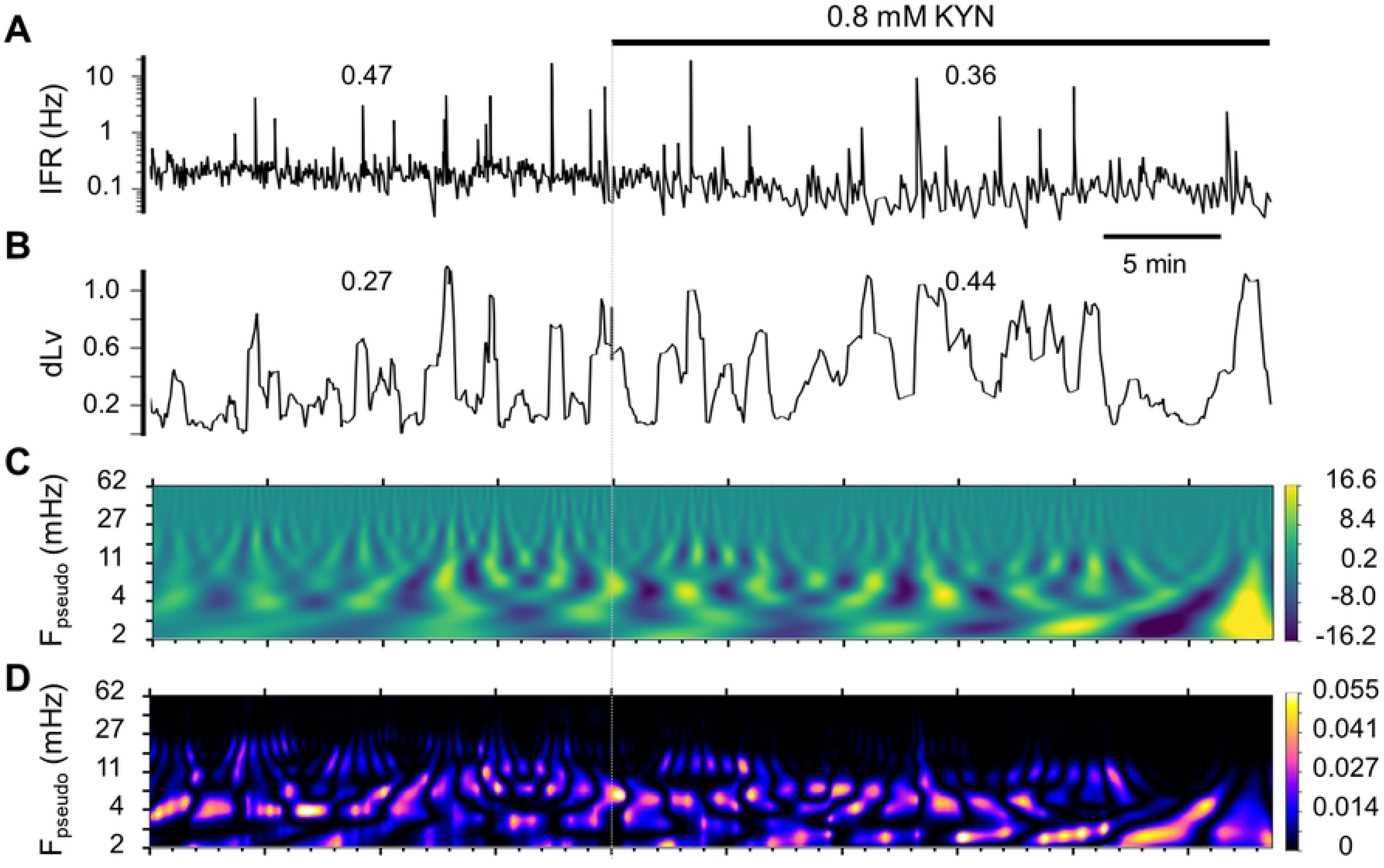
Continuous wavelet analysis of dLv oscillations along the time course of KYN application showing a quasiperiodic change in the oscillatory patterns. Numerical values on top of the traces in **A and B** are mIFR and sLv, respectively. **A:** KYN application causes a gradual decrease of IFR. **B:** KYN application alters the dLv oscillatory pattern, showing a quasiperiodicity with extended cycle length. **C:** A color map representation of the coefficients of wavelet transform (Cwt) along the frequency and time axes. F_pseudo_, pseudo-frequency. Quasiperiodic appearance of intense Cwt was apparent at ∼4‒11 mHz under control conditions, which was substantially altered by KYN application. **D:** A wavelet energy scalogram (Wes) showing the probability density of energy (PD_E_) along the frequency and time axes. KYN application increased PD_E_ at ∼ 2 mHz and decreased PD_E_ at ∼11‒27 mHz.

**Fig 6.**
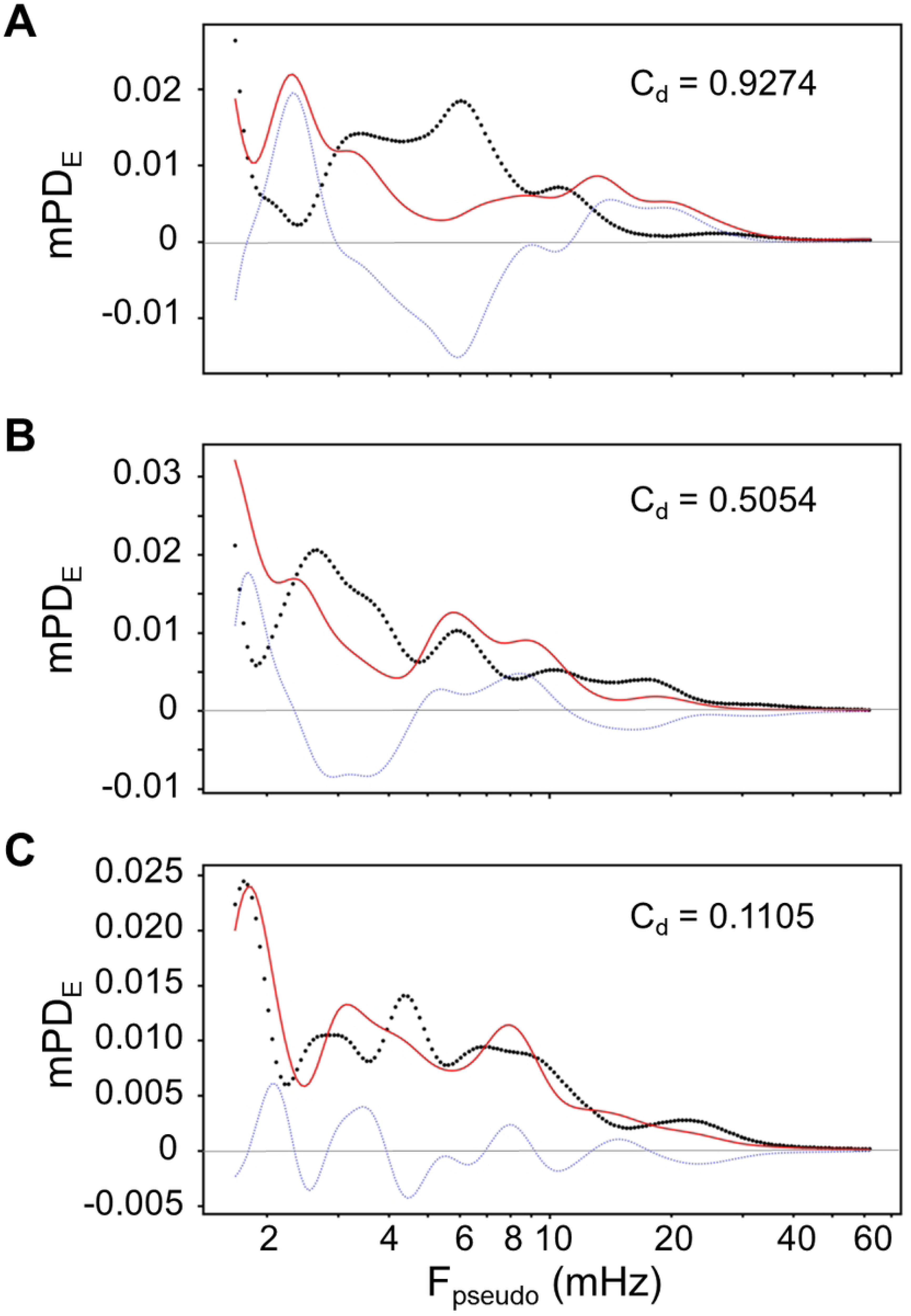
Plots of the mean of PD_E_ (mPD_E_) versus the F_pseudo_ showing the energy profiles of the dLv oscillations. Changes of the energy profiles were induced by application of KYN **(A)**, CNQX **(B)** or APV **(C)**. In each panel, the thick black dotted line and the red line represent the data acquired under control conditions and after antagonist application, respectively; the thin blue dotted line is acquired by subtracting the mPD_E_ under control from the mPD_E_ after antagonist application. C_d_, coefficient of deviation, estimates the dissimilarity between the curves acquired under control conditions and after antagonist application. Compared to panels **B** and **C**, panel **A** had the highest C_d_, showing that antagonist application produced a prominent change of the energy profiles of the dLv oscillations.

### Power-based and rate-dependent modulation of dLv rhythmicity

In signal processing, mPD_E_ could be taken as an estimate of the energy levels of the dLv rhythmogenesis under certain experimental conditions. The power of dLv oscillating at a frequency is defined as P_f_, which can be easily calculated as the product of mPD_E_ and its corresponding frequency (i.e., F_pseudo_, a pseudo-frequency component acquired by continuous wavelet analysis). P_f_ is measured in energy per unit time, describing the power of dLv with rhythmic oscillations when F_pseudo_ = ‘f’. Thus, P_f_ could estimate the power of the rhythmic signals at the designated frequencies. Fig 7 demonstrates the conversion of mPD_E_ curves into P_f_ curves, showing plots of P_f_ in the frequency-domain. Therein, the maximal P_f_ peaked at 15.11 mHz of 0.0996 in magnitudes under control conditions and at 7.94 mHz of 0.0736 in magnitudes after PIC application. To reveal the dominant dLv rhythmicity at the population level, the data obtained from the 84 fibers were pooled. Under control conditions an average of the maximal P_f_ frequency was 7.44 ± 0.54 mHz (range: 1.67‒20.76 mHz, *n* = 84). For all testing groups, the drug applications did not significantly change their maximal P_f_ peak locations in the frequency-domain. However, the magnitudes of the maximal P_f_ peaks were significantly reduced by application of APV (Control vs. APV: 0.1105 ± 0.0073 vs. 0.0816 ± 0.0030, *P* < 0.001; *n* = 22) or CNQX (Control vs. CNQX: 0.1105 ± 0.0072 vs. 0.0921 ± 0.0051, *P* < 0.05; *n* = 17), indicating a form of power-based modulation of the dominant dLv rhythmicity via the neural network activity mediated by NMDA or non-NMDA receptors.

**Fig 7.**
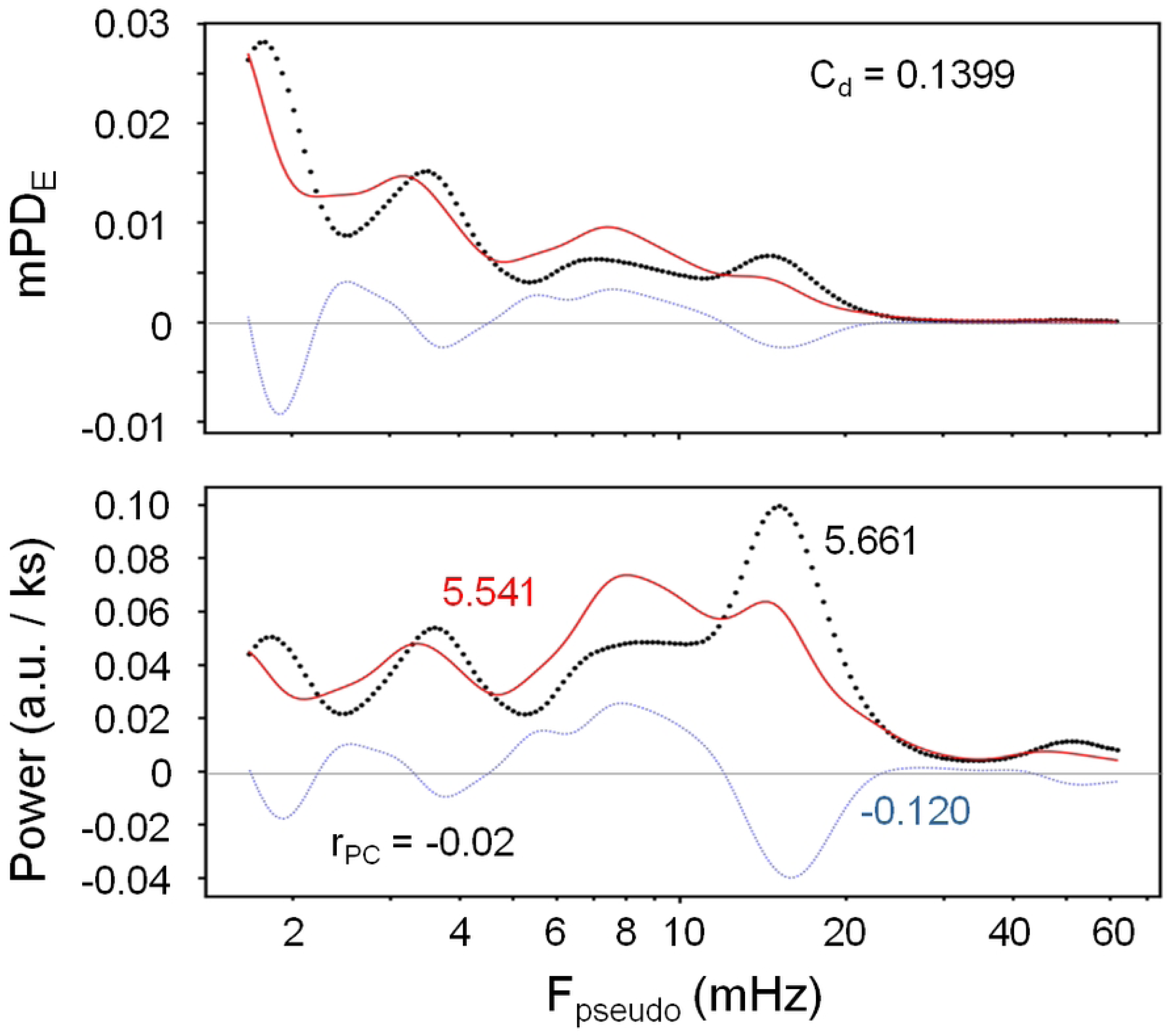
An example showing a conversion of mPD_E_ into power. In both panels, the thick black-dotted lines are acquired under control conditions, the solid red lines are acquired after PIC application, and the thin blue-dotted lines are acquired by subtracting the control curve from the PIC curve. By multiplication of mPD_E_ with their corresponding frequencies, the lower panel shows the power curves. For each power curve, as color-coded, the numerical values are the sum of the power spanning across all the frequency bands. The ratio of the power change (r_PC_) was - 0.02, as calculated by -0.12 / 5.661. a.u., arbitrary units of energy.

Power-based modulation of the dominant dLV rhythmicity was not observed in the testing groups of KYN and PIC. Although application of KYN or PIC could reshape the maximal P_f_ in individual fibers, both tests did not significantly change either the averaged frequency or the averaged magnitudes of the maximal P_f_ at the population level. However, in both groups, the drug application caused a spiking rate-dependent shift of the peak locations of the maximal P_f_ in the frequency-domain. Fig 8 shows the scatter plots of the quotients of the drug-induced change in maximal P_f_ frequency versus the quotients of drug-induced change in mISI, showing the data distributed in a decaying power-law manner. Because mISI is inverse to AFR, those data showing a decreased and increased firing after KYN or PIC application were distributed to the right and the left of the y-axis with q_mISI_ >1and <1, respectively. The decaying power-law relationship as shown in Fig 8 indicated that an elevated firing activity was accompanied with a shift of the dominant dLv rhythm to a higher frequency range, and a lower firing activity with a shift to a lower frequency range. By contrast, the abovementioned decay power-law relationship was not observed in the testing groups of APV or CNQX. These findings indicated a form of spiking rate-dependent frequency modulation of the dominant dLv rhythmicity via the neural network activity mediated by a broad spectrum of ionotropic glutamate receptors or GABA_A_ receptors.

**Fig 8.**
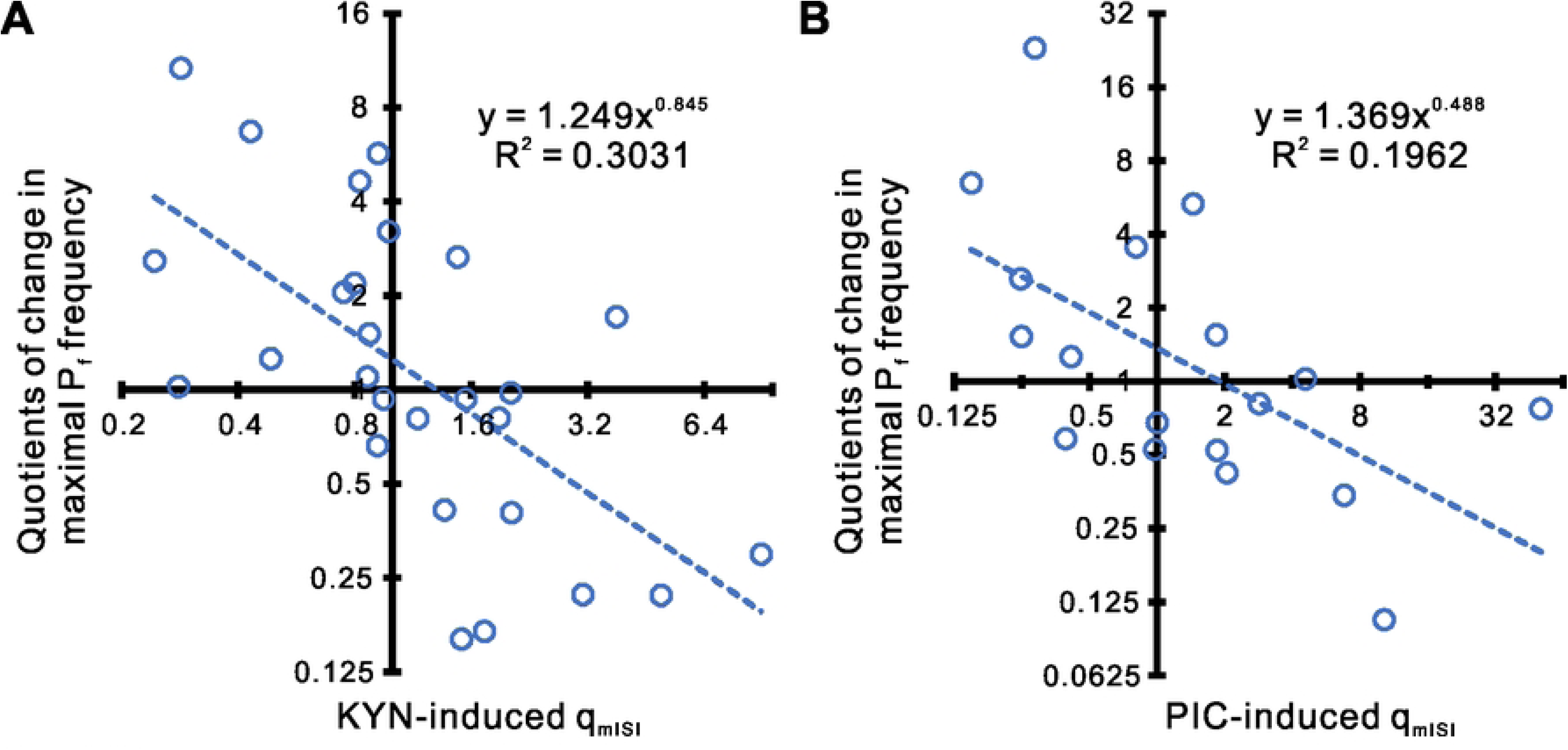
Decay power-law relationship between the maximal P_f_ shifting and the mISI change after application of KYN or PIC. In a log-log scale, plotting the quotients of the antagonist-induced shift in the frequency locations of the maximal P_f_ versus the quotients of the mISI change shows a linear data distribution. The data were significantly regressed by the power-law functions with *P* < 0.001 in **A** (*n* = 27) and *P* < 0.05 in **B** (*n* = 18).

### Concomitant changes in dLv oscillatory power and firing activity

Abovementioned results clearly demonstrate that antagonist application alters the neural network operation and causes a substantial change in the frequency- or the magnitude-components of dLv oscillations. Because P_f_ is the dLv oscillatory power at a frequency, a simple summation of all the P_f_ across the whole frequency bands of interest (i.e., ΣP_f_) can estimate the total power required for dLV rhythmogenesis. For instance, as shown in Fig 7, ΣP_f_ was 5.661 under control and was reduced to 5.541 after PIC application, yielding a ratio of power change (r_pc_) -0.02 (i.e., r_pc_ = (5.541-5.661) / 5.661). Such an antagonist-induced change in ΣP_f_ suggested that ΣP_f_ can be taken as an energy expenditure of its underlying neural network and correlate with the neural activity. To address this point, whether a drug-induced change of the dLv oscillatory power was concomitant with a change in its firing activity was explored. Fig 9 shows 3 selected examples to demonstrate that a positive or negative value of r_pc_ well describes the direction of changes in firing activity, i.e., a positive r_pc_ entails an increased firing activity and a negative r_pc_ a decreased firing. To further recapitulate the mathematical relationship between the r_pc_ and the change of firing activity, whether the q_mISI_ could be described as a function of the r_pc_ was further explored. Three aspects of constraints were considered in choosing a mathematical model. First, the extent of mISI shortening, which might result from an enormous excitation of the neural circuit operation due to a highly elevated r_pc_, would be limited by the refractory period of the action potential generation and reach a plateau. Second, the extent of mISI lengthening, which might result from an inhibition of the neural circuit operation due to a negative r_pc_, would be limited by an intrinsic firing property (e.g., pacemaker-like activity) and also reach a plateau. Third, the dependence of the drug-induced change of mISI on r_pc_ might not be linear. Taken these constraints together, a negative sigmoid model was chosen for curve fitting. Data were pooled from all the 84 fiber responses. Fig 10 shows a semi-log plot of q_mISI_ versus r_pc_ and demonstrates that the data distribution is well described by a negative sigmoid curve. Notice that the data with log(q_mISI_) > 0 are those with decreased firing, which have an increased mISI after an antagonist application and yield a q_mISI_ > 1. Conversely, the data with log(q_mISI_) < 0 are those with increased firing, which have a decreased mISI after an antagonist application and yield a q_mISI_ < 1. The acquired equation predicts that a 100-fold increase of ΣP_f_ (i.e., r_pc_ = 100) would yield a q_mISI_ of 0.19 (i.e., a shortening of mISI to 0.19 of its control value) and a complete removal of ΣP_f_ after drug application (i.e., given r_pc_ = -1) would yield a q_mISI_ of 99.86 (i.e., ∼100-fold lengthening of mISI to its control value). Thus, the heterogeneity in drug-induced firing responses, showing a wide distribution of q_mISI_ with values deviated from 1 (i.e., log(q_mISI_) = 0), is explained by r_pc_. This result supports the view that the dynamic of the neural activity is manifested by dLv oscillations and the dLv oscillatory powers reflect the activity states of underlying neural network operation.

**Fig 9.**
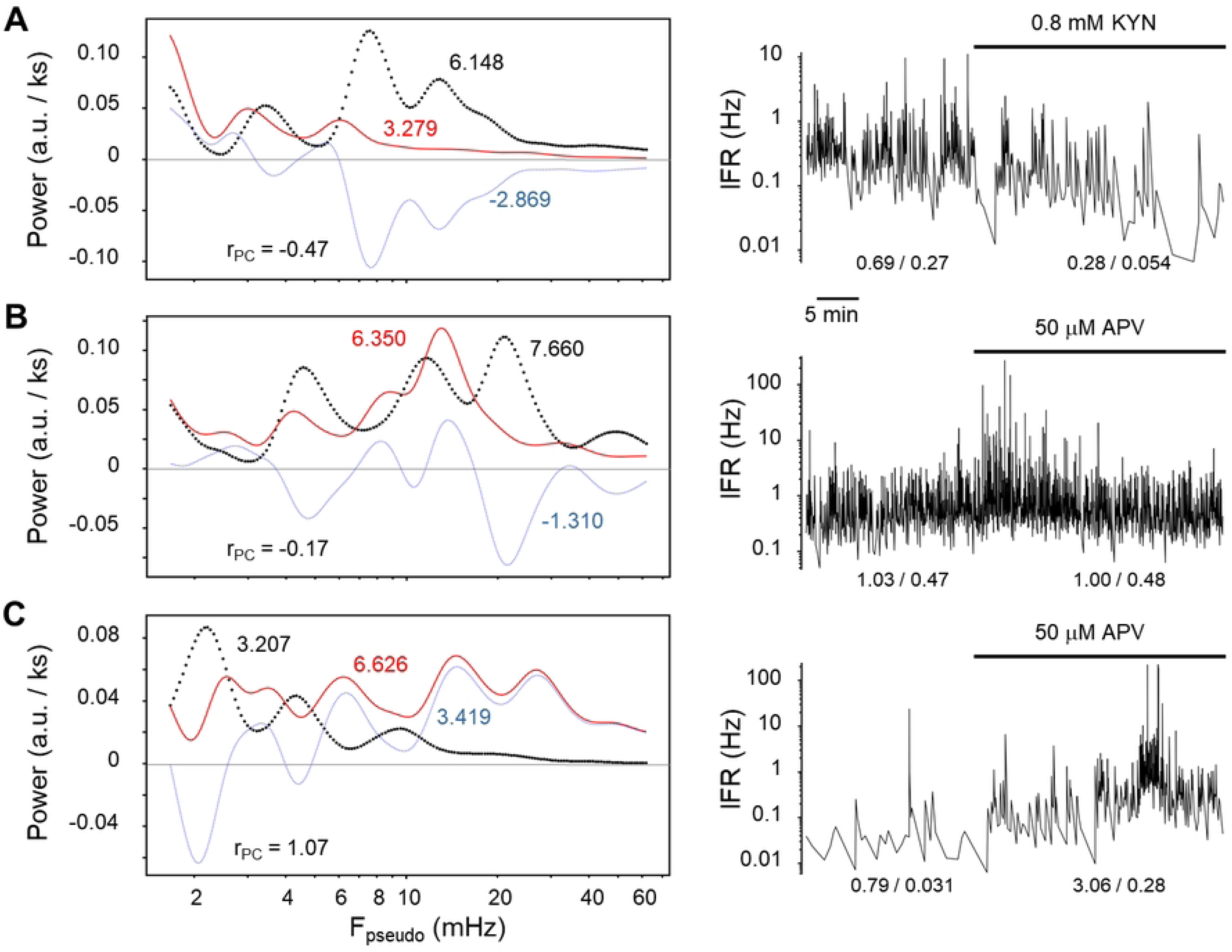
Three selected examples showing that the extents of antagonist-induced change in firing activity are correlated with r_PC_. The lines and the numerical values in the left panels were as explained in **Fig 9**. The right panels show the time course of IFR; the numerical values at the bottom are mIFR / AFR. From **A** to **C**, an increment of r_PC_ from negative to positive values is accompanied with an antagonist-induced firing response from inhibitory to excitatory.

**Fig 10.**
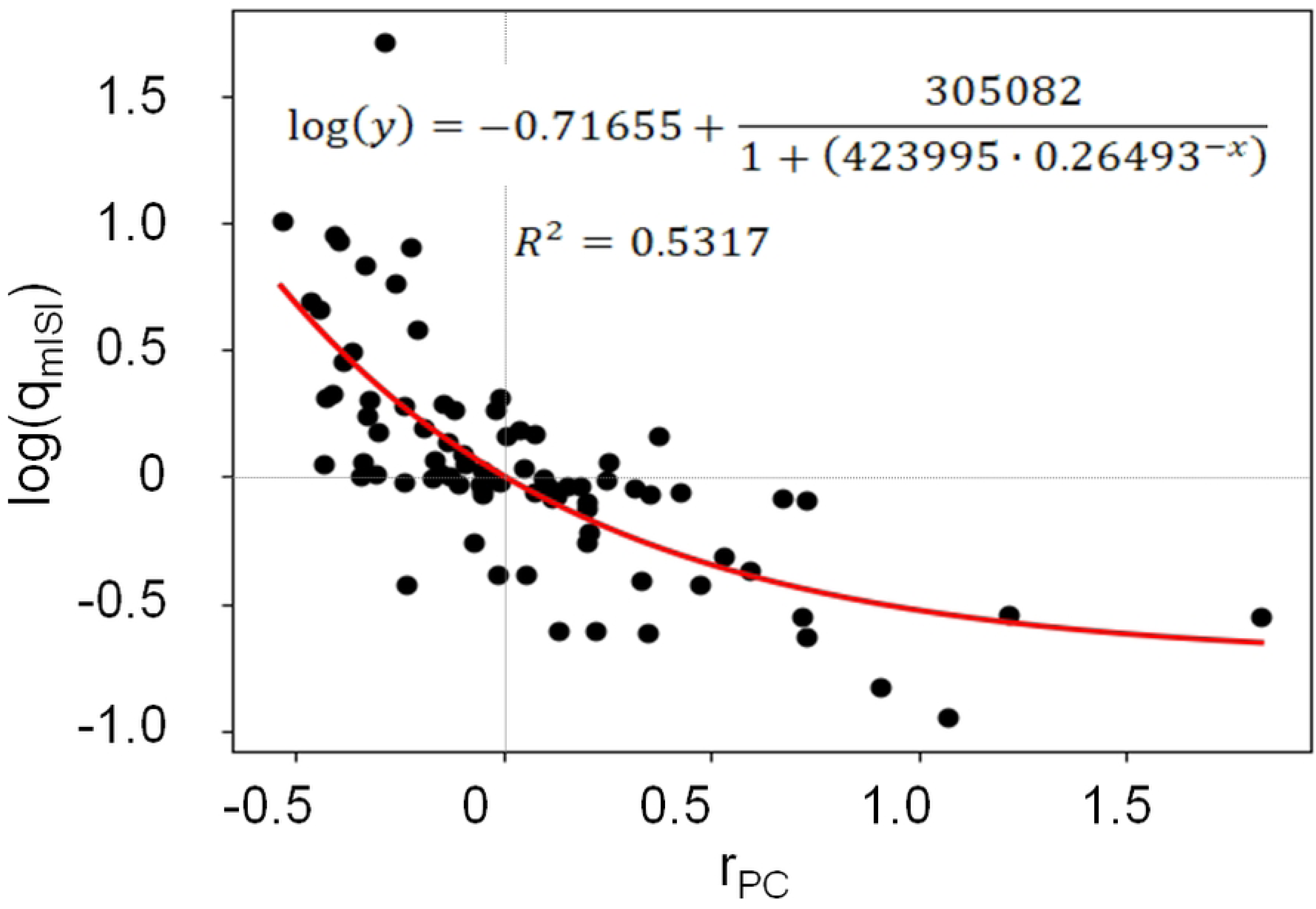
A scattered plot of log(q_mISI_) versus r_PC_ is fitted by a negative sigmoid growth curve. Data were pooled from all the experiments (*n* = 84). The regression was significant with *P* < 0.0001. The vertical dashed line setting at r_PC_ = 0 and the horizontal dashed lines setting at log(q_mISI_) = 0 intersect with the fitted curve at the points when q_mISI_ = 1.0069 and when r_PC_ = 0.0031, respectively. These intersecting points approximate a theoretically null point where r_PC_ = 0 and q_mISI_ = 1 (i.e., log(q_mISI_) = 0).

## Discussion

This study used Lv as a metric to quantify sympathetic spiking patterns. The dynamic components of the spiking patterns were revealed by dLv oscillations, which registered a dominant rhythm at 7.44 ± 0.54 mHz. Intraspinal neurotransmitter activities mediated by ionotropic glutamate receptors and GABA_A_ receptors heterogeneously modulate the frequency and/or the magnitude of dLv oscillations, and thus, participate in tuning the spiking patterns. Most intriguingly, as dLv power change well correlates with the dynamic of neural responses, dLv oscillations can be taken as a functional display of the neural network activity. As it is discussed below, the rhythmicity embedded in dLv oscillations seems to format the central visceral commands and mirror the functional operations of their peripheral target organs.

### A quantitative expression of the neural network activity by Lv

The dynamic components of Lv as revealed by the dLv oscillations along the time course were largely changed after interrupting the endogenous neural network activity mediated by ionotropic glutamate receptors or GABA_A_ receptors. This was indicated by 68/84 fibers with their dLv significantly changed by the drug application. Also, using C_d_ as a quantitative measure of the antagonist-induced change in mPD_E_ profiles, all the testing groups had an averaged C_d_ of values ∼0.5, which was significantly different from a hypothetical C_d_ of value 0 if an mPD_E_ profile was not altered by the antagonist application. These findings clearly indicate that dLv oscillations were driven by a neural network activity.

Heterogeneity in drug-induced firing responses is basal activity-dependent [23, 26–28]. In other words, an antagonist application tends to suppress those sympathetic fibers with higher basal firing rates and boost those with lower ones. Heterogeneity in drug-induced dLv power changes was also observed in this study. Intriguingly, change in dLv oscillatory powers correlated with the dynamic of heterogeneous firing responses. Because dLv power could be taken as a quantitative measure of the energy underlying neural network operations, an increase or decrease of dLv power unveiled an elevation or reduction of the energy levels of the network operation, which subsequently leads to a concomitant increase or decrease of neural firing. Thus, from the view point of the energy expenditure required for neural firing, the magnitudes of dLv oscillatory power can be used to gauge the activity states of neural network operations.

In earlier reports [20, 29], Lv is believed to reflect the neuronal excitability. This study provides evidence and demonstrates that a substantial change of dLv is often elicited by interrupting endogenous amino acid neurotransmitter activities, confirming that Lv indeed is a metric reflecting the outcome of synaptic integration.

### Conveyance of central visceral commands by dLv oscillations

How the central nervous system configures the visceral commands in a format that can effectively regulate peripheral organs was not clear. Many in vivo studies demonstrated that the sympathetic nerves convey spiking activities in ranges of Hz. In chloralose-anesthetized chronic spinal cats, 1-6 Hz slow wave activity was observed in the splanchnic sympathetic nerves [30]. In Sprague-Dawley rats that were initially anesthetized by pentobarbital and with supplemental anesthesia by α-chloralose, the postganglionic sympathetic nerves innervating the tail artery have an ongoing activity at 0.4‒5.9 Hz [31]. In pentobarbital-anesthetized Wistar rats’, the rate of spontaneous single-fiber activity recorded in the postganglionic sympathetic nerves innervating the tail artery is 0.23‒2.6 Hz [32]. Under the current in vitro control conditions, spontaneously active sympathetic preganglionic neurons have AFR of 0.08‒2.32 Hz with a population mean of AFR at 0.72 ± 0.04 Hz [13]. As the spiking rates as we observed here were largely comparable to those from in vivo preparations, the rhythmicity embedded in dLV oscillations as we observed here could mimic those under in vivo experimental conditions.

Abiding to a rate-code theory for information processing, sympathetic effectors should respond to the firing rates being conveyed in the nerve traffics up to several Hz. However, sympathetic targeted organs have long been known as a low-pass filter and fail to respond timely to the rates of the sympathetic spiking. Several classical studies have addressed this point and show that the effectiveness of transferring sympathetic motor commands to their target organs is limited to a rhythmic frequency range < tens of mHz [15–18, 33]. In this study, the temporal order of ISIs formulated the spiking patterns, unveiling dLv oscillations at 7.44 ± 0.54 mHz. This slow rhythm registered only tenth to hundredth of the spiking rates. Resonance of the dLv rhythmicity with the sympathetic effectors renders dLv oscillations a realistic coding format that conveys central visceral commands.

In line with the notion that dLv oscillations encode central visceral commands, endogenous receptor activities modulate the commands in frequency- and/or power-based manners. Because dLv oscillations address the dynamics of spiking patterns, its oscillatory power refers to an extent of pattern change and its rhythmic frequency refers to a periodic pattern change in time. Many sympathetic effectors are hollow organs lining with smooth muscle layers. Conveyance of the rhythmic commands to the smooth muscles would produce a dynamic visceral motor pattern that varied in both extents and periodicities and lead to a quasiperiodic contractile movement in forms that varied from tonic constrictions when dLv = 0 to epileptic spasms when dLV >1.

It should be recalled that most visceral organs receive central commands via a synaptic relay in the sympathetic ganglions. Ample evidence supports the notion that the central visceral commands conveying by the preganglionic fibers are faithfully carried out by the sympathetic ganglionic neurons (SGNs). First, SGNs receive strong excitatory synaptic inputs from the preganglionic fibers [34–37]. Second, some SGNs are activated by electrical stimulation of a single preganglionic fiber [38, 39]. Third, one sympathetic preganglionic neuron innervates a subset of SGNs [38, 40]. Fourth, 10% of sympathetic preganglionic fiber pairs fire synchronously to facilitate ganglionic synaptic transmission [26]. Thus, it is likely that the central visceral commands being formatted as the dLv oscillations can also be replayed in the sympathetic ganglions and conveyed downstream to regulate their peripheral effectors.

### Frequency-matching operations of dLv rhythms with sympathetic effectors

Sympathetic effectors operate at different rhythms. For instance, in mice, the gastric rhythm is ∼7 cycles per minute (cpm) and the small intestinal rhythms vary from ∼22 to ∼27 cpm [41]. As 1 cpm equals to 16.7 mHz, the gastrointestinal motility in mice operates in a frequency ∼100‒450 mHz. In the smooth muscle cells of the monkey aorta, an actin-myosin driven process causes a temporal oscillation of elasticity at a frequency from tenth to tens of mHz [42]. Besides, mechanically stretching or chemical activation of a variety of smooth muscle cells cause intracellular Ca^2+^ oscillations at frequencies from tenth to tens of cpm [43, 44]. Most intriguingly, stimulation of the smooth muscle cells in the intrapulmonary arterioles with 100 mM KCl causes an intracellular Ca^2+^ oscillation at a frequency ∼0.5 cpm (i.e., ∼8 mHz; see Fig 5 in [45]). This is a frequency response that is very close to the dominant dLv rhythm 7.44 ± 0.54 mHz as reported here. Because Ca^2+^ oscillations could cause rhythmic constrictions of smooth muscles, these findings implicated that the sympathetic effectors operate in a feature with wide frequencies. In this study, the dominant dLv rhythms vary substantially between individual fibers. The diversity of spiking patterns between fibers and the dynamics of spiking patterns subjected to neuromodulation in individual fibers may help to lay out a scheme for broad-band frequency-matching operations with their effectors. In conclusion, the temporal structures of sympathetic spiking activity construct the central visceral commands that are manifested as a slow rhythmic change of spiking patterns, and by which, to echo the frequency response of their peripheral effectors.

## Materials and Methods

### Experimental animals

Experiments were performed using 40 neonatal Sprague-Dawley rats at age 1-6 postnatal days. All surgical and experimental procedures were approved by the Institutional Animal Care and Utilization Committee of Academia Sinica (Protocol#: RMiRaIBMSC2011081) in accordance with the Guide for the Care and Use of Laboratory Animals of the Agriculture Council of Taiwan.

### Splanchnic sympathetic nerve–thoracic spinal cord preparations *in vitro*

*En bloc* preparations retaining the splanchnic sympathetic nerve–thoracic spinal cord (T1–T12) were prepared following surgical procedures as previously described [11, 23]. Briefly, neonatal rats were rendered unconscious by hypothermia [46], followed by prompt midcollicular decerebration. During dissection, the reduced preparation was immersed in ∼4 °C artificial cerebrospinal fluid (aCSF; in mM: 128 NaCl, 3 KCl, 1.5 CaCl_2_, 1.0 MgSO_4_, 24 NaHCO_3_, 0.5 NaH_2_PO_4_, 30 D-glucose and 3 ascorbate; equilibrated with 95% O_2_-5% CO_2_). A stub of splanchnic sympathetic nerves was freed from surrounding tissues. The nerve stub was predominantly composed of sympathetic preganglionic fibers and was dissociated by 0.5% type IV collagenase (C5138, Sigma-Aldrich). During experiments, the nerve-thoracic spinal cord preparation (T1–T12) was immersed in a bath chamber containing 30 ml of freshly-oxygenated aCSF with the temperature maintained at 24.5 ± 1 °C.

### Neural recordings and signal processing

The procedures regarding simultaneous recordings of several sympathetic fiber activities and their signal processing were as previously described [24]. Briefly, analog signals obtained from the neural recordings were digitized in real time using a National Instrument-based data acquisition system (NI-PCI-6010, National Instrument, Austin, Texas) and processed using a customized LabVIEW program (version 15.0.1.1f2, National Instrument) incorporated with MATLAB scripts (version 8.5.0 The MathWorks, Inc., Natick, Massachusetts). A series of computational algorithms were used for spike sorting to acquire the sympathetic single-fiber activities. Times of spike occurrence in each fiber were automatically detected at the spike peaks. The spiking times were further processed to acquire the metrics describing the spiking activity using a series of Python-based programs (version 3.8), available in the website: https://github.com/chunkuei/LvAnalysis

### Acquisition of the metrics describing the spiking activity in rates and variations

Based on an epoch of spiking events containing *n* ISIs, the following metrics were calculated to describe the spiking activity in the contexts of rates and variations. The inverse of the means of a series of ISIs (mISI) defines the average firing rates (AFR), which are acquired by the equation:

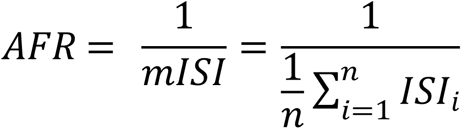

where *i* is the index of the i^th^ ISI. The i^th^ Instantaneous firing rate (IFR_i_) is defined as the inverse of ISI_i_ and the mean of a series of IFR_i_ (mIFR) is acquired by the equation:

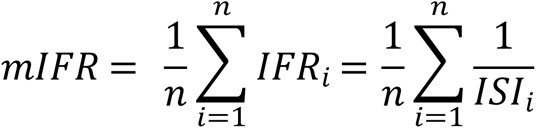

In an ideally regular firing, all ISIs are equal and AFR are equal to mIFR. Thus, in a series of spiking events with unequal ISIs, the firing rate irregularity (FR_ir_) can be estimated by the equation:

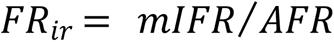

The coefficient of variation (Cv) of a series of ISIs is a conventional index to describe the variation of the spiking activity and is acquired by the equation:

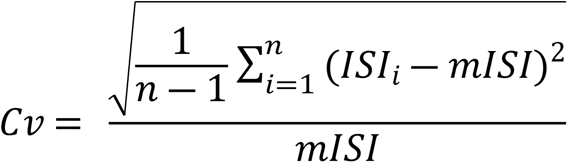

The local variation (Lv) of the spiking activity, which is relatively unaffected by the rate of the spiking activity, is calculated by the following equation [22]:

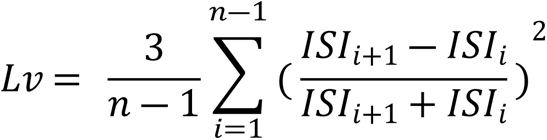

Using the above equation, based on a difference in the selections of *n*, two types of Lv were derived. One is the static Lv (sLv) acquired by taking all the ISIs under certain experimental conditions into account. The other is the dynamic Lv (dLv) acquired by iteratively calculating the Lv based on successive epochs of spiking events along the time course of experiments, which yield a series of dLv with sequential timestamps to show the Lv fluctuations. To obtain an optimal temporal resolution of the dLv, an even number of ISIs (N_ISI_), which is approximately equivalent to an epoch of 32-s of spiking activity under control conditions, is determined by the equation:

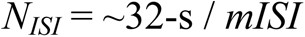

where mISI is the mean of the ISIs under control conditions. Accordingly, in a sequence of N_ISI_ + 1 spiking events, a timestamp of the acquired dLv was assigned by the middle event of the spiking sequence, i.e., the occurrence time of the (*N_ISI_*/2) +1 spiking event. In experiments with the mISI > 8-s, N_ISI_ was arbitrarily set to 4. Analyses only include experiments with mISI ≤ 50-s so as to limit the acquisition of dLv from epochs of spiking events with averaging time spans ≤ 200-s for a better temporal resolution.

### Continuous wavelet analyses of dLv oscillations

To avoid a presumption of signal stationarity, continuous wavelet transforms were used to extract the rhythmic features of dLv oscillations. Original dLv signals appeared at unequal intervals and were thus resampled by interpolation methods at 0.05-s intervals, i.e., a resampling rate of 20 Hz. A size of 160 scales covering a pseudo-frequency (F_pseudo_) range of 1.67‒61.6 mHz spaced in log-scales were defined to evaluate the coefficients of wavelet transforms (Cwt) using the Morlet wavelet as a mother function 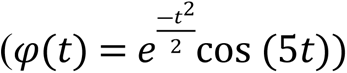. The Cwt in time-frequency domain (*Cwt(a, b)*) are computed by the following equation [47]:

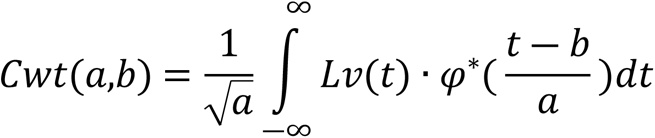

where the asterisk denotes the complex conjugate of the Morlet wavelet 𝜑(*t*), *a* is a scale factor inversely proportional to the F_pseudo_, and *b* is a translation factor indicating the time points in the time domain. Based on the central frequency of the Morlet wavelet (i.e., 0.8125 Hz at 1-Hz sampling rate), the F_pseudo_ with a scale factor ‘*a*’ being defined as *f_a_* is calculated by the equation:

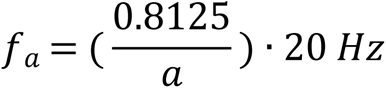

where 20 Hz is the resampling rate of the dLv. In practice, *Cwt(a, b)* were simply acquired by employing a Python wavelet module.

Sinusoidal-like oscillations of the *Cwt(a, b)* along the time course of the experiments yielded a mean coefficient of 0 across the time domain. Thus, the square of *Cwt(a, b)* was taken as a wavelet energy and was used to extract the power components in the frequency domain. First, a normalized wavelet energy scalogram (Wes) was acquired by the following equation [48]:

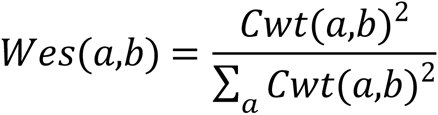

The sum of the normalized *Wes(a, b)* across all the scale factors at any time points equals to 1. Thus, *Wes(a, b)* shows the probability density distribution of the energy of *Cwt(a, b)* in a time-frequency domain and displays the relative intensity of the rhythmic signal at a F_pseudo._ Each value in *Wes(a, b)* at a time point is then defined as the probability density of energy (PD_E_). Second, an average of *Wes(a, b)* across a selected period of time under certain experimental conditions reveals the energy-frequency features and defines the means of the PD_E_ (mPD_E_), comprising 160 values with each representing the relative intensity of the signals at a F_pseudo_. Third, because power is the product of energy and frequency, P_f_ was then defined as the power at a F_pseudo_, expressed in arbitrary units of energy per unit time, and was acquired by the equation:

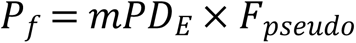

P_f_ represents the driving power of the signals at a frequency when F_pseudo_ = ‘*f*’. The whole spectra of F_pseudo_ ranging from 1.67 to 61.6 mHz comprised 160 scales. Thus, a sum of all the 160 P_f_ (i.e., ΣP_f_) was used to estimate the total power of dLv oscillations under certain experimental conditions.

### Drugs and drug applications

Kynurenic acid (KYN, a broad-spectrum ionotropic glutamate receptor blocker), 6-cyano-7-nitroquinoxaline-2,3-dione-disodium salt (CNQX, a non-NMDA receptor blocker), and picrotoxin (PIC, a GABA_A_ receptor blocker) were purchased from Sigma-Aldrich, and L-2-amino-5-phosphonopentanoic acid (APV, an NMDA receptor blocker) from Tocris. APV, CNQX, and KYN were dissolved in water and PIC in DMSO to prepare concentrated stock solutions. A final concentration of 0.8 mM KYN, 20 μM CNQX, 50 μM APV, or 100 μM PIC was achieved by adding an aliquot of the stock solution directly to the bath chamber. An elapse of 5 min after drug applications was allowed for equilibration. For each experiment, the neural recording in a drug test consists of 20-min of control followed by 30-min after a drug application. Unless otherwise mentioned, the drug-induced responses were evaluated by comparing the firing activities that occurred in a 20-min epoch prior to a drug application with those in the last 20-min epoch after the drug application.

### Data analysis

The values of the metrics, including AFR, mIFR, FR_ir_, Cv, sLv, and dLv means, were automatically acquired from epochs of data prior to and after drug applications using Python-based programs as abovementioned. For individual fibers, series of the dLv values acquired along the time course of the drug tests were used to calculate their dLv means. Our previous studies demonstrate that a drug administration often exerts a heterogeneous, power-law modulation of the sympathetic firing [23, 26–28]. To further explore if the drug administrations in this study also changed the temporal structure of firing in a similar manner, the data resulted from the drug-induced change in some of the metrics that described the spiking patterns (e.g., sLv) were examined in a log-log plot and further regressed by a power function using Microsoft Office Excel (version 12.0.6787.5000):

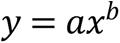

where *x* and *y* are the metrics of interest, *a* is the intercept, and the exponent *b* is the slope.

In wavelet analysis, the largest scale factor chosen in the continuous wavelet transforms is equivalent to a period of 600-s, which causes an edge problem of acquired *Cwt(a, b)* or *Wes(a, b)* in a cone shape with a time-base up to 300-s [49]. To avoid the edge problems and minimize the data loss, the mPD_E_ under control conditions was obtained from a 10-min epoch of *Wes(a, b)* starting from 5 min after the neural recording and ending at 5 min prior to a drug application, whereas the mPD_E_ during a drug test was obtained from an epoch of *Wes(a, b)* starting from 5 min after a drug application and extending indefinitely till a time point 15-s prior to the end point of *Wes(a, b).* Drug-induced change of mPD_E_ was evaluated by the coefficient of deviation (C_d_), which was calculated by the equation:

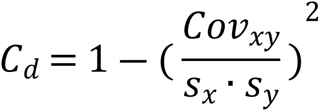

where *Cov* is the covariance, *s* the standard deviation, *x* the mPD_E_ under control, and *y* the mPD_E_ after the drug application. For each fiber activity, the significance of the drug-induced change in the dLv was evaluated by Student’s *t*-test, wherein the means calculated from series of the dLv under control conditions and after a drug application were compared. The quotients of the drug-induced change in a parametrical value (e.g., the quotient of mISI, q_mISI_) were calculated by dividing the parametrical value under control conditions from the parametrical value after a drug application. A ratio of power change (r_pc_) was used to estimate a drug-induced change in dLv oscillatory powers, which were calculated by the following equation:

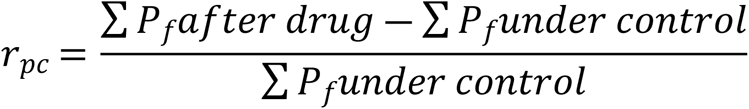

As it would be explained later, using a Python-based program that was written with algorithms of the least-squares methods, the mathematical relationship between q_mISI_ and r_pc_ was explored using a negative sigmoid growth model:

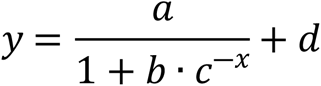

where a‒d are the parametrical values, x the r_pc_, and y the log(q_mISI_).

In general, Student’s *t*-test was used to evaluate a drug-induced effect on the metrics. Unless otherwise mentioned, values are expressed as means ± SEM. *P*-values < 0.05 are considered significant.

## Acknowledgments

The original data were collected in IBMS, Academia Sinica. Writing the Python-based programs for data analysis and preparation of the manuscript were complete when C.-K. S. accepted a tentative lectureship in GBTU . The authors are grateful to Ms. Ya-Ting Chang and Ms.Yu-Pei Fan for excellent technical assistance, and Dr. Chok-Yung Chai for general supports.

## Author Contributions

**Conceptualization:** Chun-Kuei Su, Chiu-Ming Ho.

**Formal analysis:** Chun-Kuei Su.

**Investigation:** Chun-Kuei Su.

**Software:** Chun-Kuei Su.

**Validation:** Chun-Kuei Su, Chiu-Ming Ho.

**Visualization:** Chun-Kuei Su.

**Writing – original draft:** Chun-Kuei Su.

**Writing – review & editing:** Chun-Kuei Su, Chiu-Ming Ho.

